# Evaluating Paraspinal Muscle Response and Compensation via Musculoskeletal Modeling in Spinal Stenosis Surgeries

**DOI:** 10.1101/2024.03.17.585440

**Authors:** Ryan Jones, Yogesh Kumaran, Adit Padgaonkar, Brett Hoffman, Kyle Behrens, Hossein Elgafy, Sudharshan Tripathi, Vijay K. Goel

## Abstract

**Introduction:** Lumbar spinal stenosis is a common cause of lower back pain and weakness in elderly patients. The gold standard treatment for this is lumbar laminectomy which involves widespread muscle damage to the multifidus, a complete loss of the posterior tension band which contains the supraspinous and interspinous ligaments. However, in recent years minimally invasive techniques such as bilateral and unilateral laminotomy have become more popular and are showing efficacy in the decompression of spinal stenosis. Due to its minimally invasive approach, the muscle retraction required for laminotomy is less intensive than that required for laminectomy. The overall body of literature on the surgical treatment of spinal stenosis is sparse in its interrogation of the biomechanical outcomes of these techniques and to our knowledge, there are no current publications that incorporate muscle forces.

**Methods:** A previously validated thoracolumbar ribcage finite element (FE) model was used for this study. Three different surgeries, traditional laminectomy, unilateral and bilateral midline sparing approaches at L4-L5 segment were simulated by removing the spinous process, supraspinous, and interspinous ligaments. The segmental range of motion (ROM) for all models were acquired and input into a musculoskeletal modelling software to calculate muscle forces.

**Results:** Unilateral and bilateral laminotomy showed similar muscle forces for every muscle group in both flexion and extension motion. While comparing the muscle forces in laminotomy to the laminectomy in extension motion displayed an increase in Iliocostalis lumborum (IL) by 12 % and multifidus (MF) by 16% and decrease in transverse abdominus (TA) by 138% and erector spine (ES) by 12%. For flexion, there was an increase in IL by 35%, and MF by 12%.

**Conclusion:** Our results highlight that laminectomy, which involves the removal of paraspinal muscles and posterior ligamentous structures to relieve stenosis, can lead to increased instability and necessitate muscle compensation, particularly in adjacent and thoracic spine segments. Conversely, midline sparing approaches such as laminotomies, are associated with decreased muscle compensation across spinal segments and enhanced stability.

## Background

Lumbar spinal stenosis remains a common cause of lower back pain and weakness especially in elderly patients, with the prevalence expected to climb to over eighteen million in the next decade [1]. While conservative treatments are first line, lumbar laminectomy has long been the gold standard of surgical intervention. The lumbar laminectomy is an invasive procedure which seeks to address pathogenic spinal stenosis by removing the vertebral lamina in its entirety. Despite being the gold standard treatment, the laminectomy procedure is one that involves widespread muscle damage to the multifidus muscles at the surgical site. Furthermore, the laminectomy procedure involves a complete loss of the posterior tension band which contains both the supraspinous and interspinous ligaments [2, 3]. This muscle damage along with the loss of the tension band may provoke spinal instability and need for revision surgery. The muscle and tendon damage incurred during a lumbar laminectomy can lead to complications, especially back pain, with a 2015 study reporting that persistent back pain accounted for 54.17% of all post-operative complications [4].

In recent years, minimally invasive techniques such as bilateral and unilateral laminotomy have become more popular and there is a growing body of literature demonstrating their efficacy in the decompression of spinal stenosis [5]. This procedure involves a minimally invasive incision followed by muscle retraction and subsequent removal of the minimum amount of lamina which the surgeon deems adequate to relieve nerve root compression. Due to its minimally invasive approach, the muscle retraction required for laminotomy is less intensive than that required for laminectomy. This is important given the documented connection between muscle retraction and post-operative back pain and disability [6]. The laminotomy procedure provides the benefit of sparing the posterior tension band which is composed of the supraspinous and interspinous ligaments as well as the multifidus muscles that are destroyed during a lumbar laminectomy [2, 7]. The midline sparing laminotomy technique involves preservation of these posterior ligaments and multifidus muscles when performed both bilaterally and unilaterally. Our previous study showed that increasing muscle damage may lead to an increased amount of stress on the adjacent segment [8].

The overall body of literature on the surgical treatment of spinal stenosis is sparse in its interrogation of the biomechanical outcomes of these techniques and to our knowledge, there are no current publications that incorporate muscle forces. We hypothesize that since a laminotomy preserves the multifidus muscles from the spinal processes that there will be minimal compensation from the muscles of the back as well as the anterior muscles, thus lowering stresses at the surgical and adjacent segment sites compared to the traditional laminectomy procedure. Furthermore, we hypothesize a laminotomy will result in similar biomechanics and lower stress between levels occurring when compared to a laminectomy. To test this hypothesis, we will be utilizing finite element simulated spinal range of motion (ROM) following these procedures which will be used as an input into a musculoskeletal model to compare the muscles forces produced by different muscles of the back following unilateral and bilateral laminotomy compared to a laminectomy.

## Methods

### Finite Element Model Creation

A non-linear, ligamentous finite element model was developed using the CT scans of a healthy male, adult spine with no abnormalities. This model contained a ribcage, thoracolumbar spine, and pelvis that was fixed at the ischial tuberosity [8]. The 3D geometry was generated from 1-mm slices of CT scans using MIMICS software (Materialise, Inc., Leuven, Belgium). Following 3D reconstruction, stereolithography geometry was imported into Geomagic Studio (Raindrop Geomagic Inc., Research Triangle Park, North Carolina, USA) to smooth surfaces and generate grid patterns for meshing. Three-dimensional geometry was meshed using IA-FE Mesh (University of Iowa, Iowa City, Iowa, USA) and Hypermesh (Altair Engineering, Inc., Troy, Michigan, USA). Assembly of this model was performed in Abaqus 2019 (Simulia, Providence, Rhode Island, USA).

Vertebral bodies were modelled as cortical bone with a shell of 0.5mm thickness containing a core of cancellous bone [9]. Cortical and cancellous bone were modelled as linear elastic, isotropic material. Intervertebral discs composed of annulus fibrosa and nucleus pulposa were modelled. The annulus was modelled as a composite solid with alternation ±30lJ collagen fibers using REBAR elements with “no compression” properties. The nucleus was simulated as a linear elastic material [8–12]. Facet joints were modelled using 3D gap elements with an initial defined clearance of 0.5mm. Ligamentous structures were simulated as hypoelastic materials with “tension only” properties. Physiological loading was simulated using a follower load based on the upper body mass at different vertebral levels, similar to a previous study [13]. This was modelled as wire connectors attached to the left and right side of the vertebral body and following the curvature of the spine [8, 12, 14]. Material properties of the FE model were derived from the literature and are detailed in Table 1 [8–11].

**Table 1:**
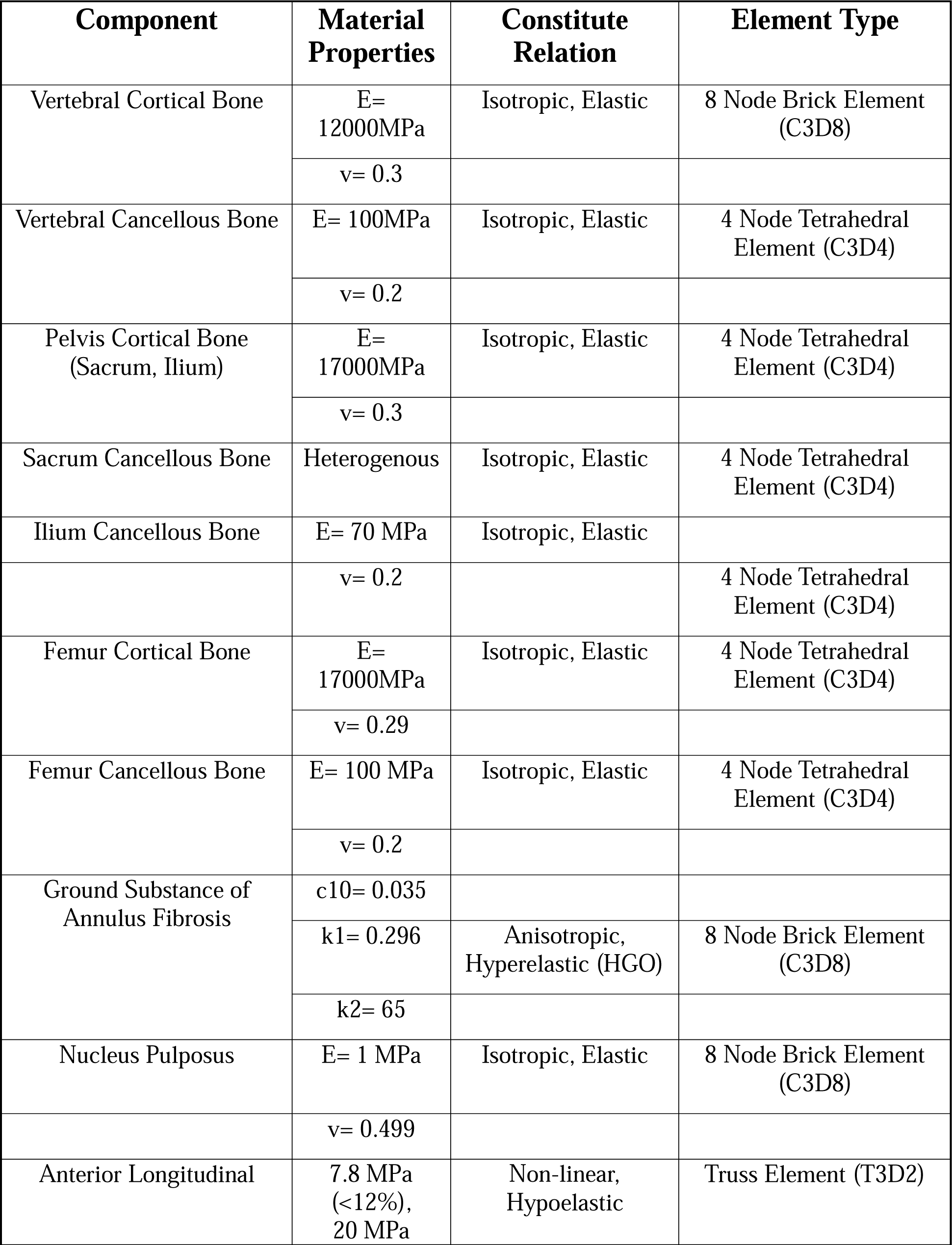
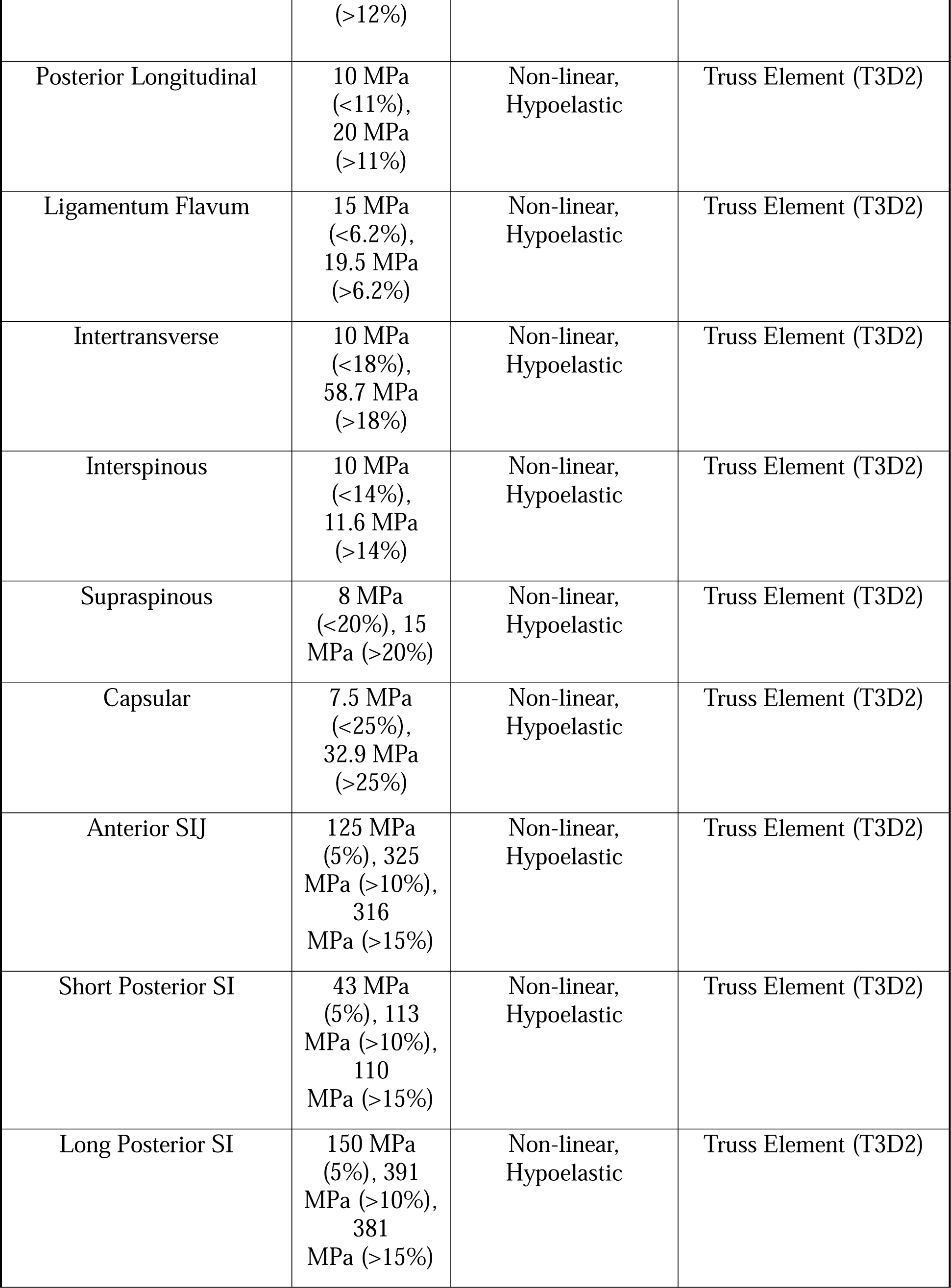
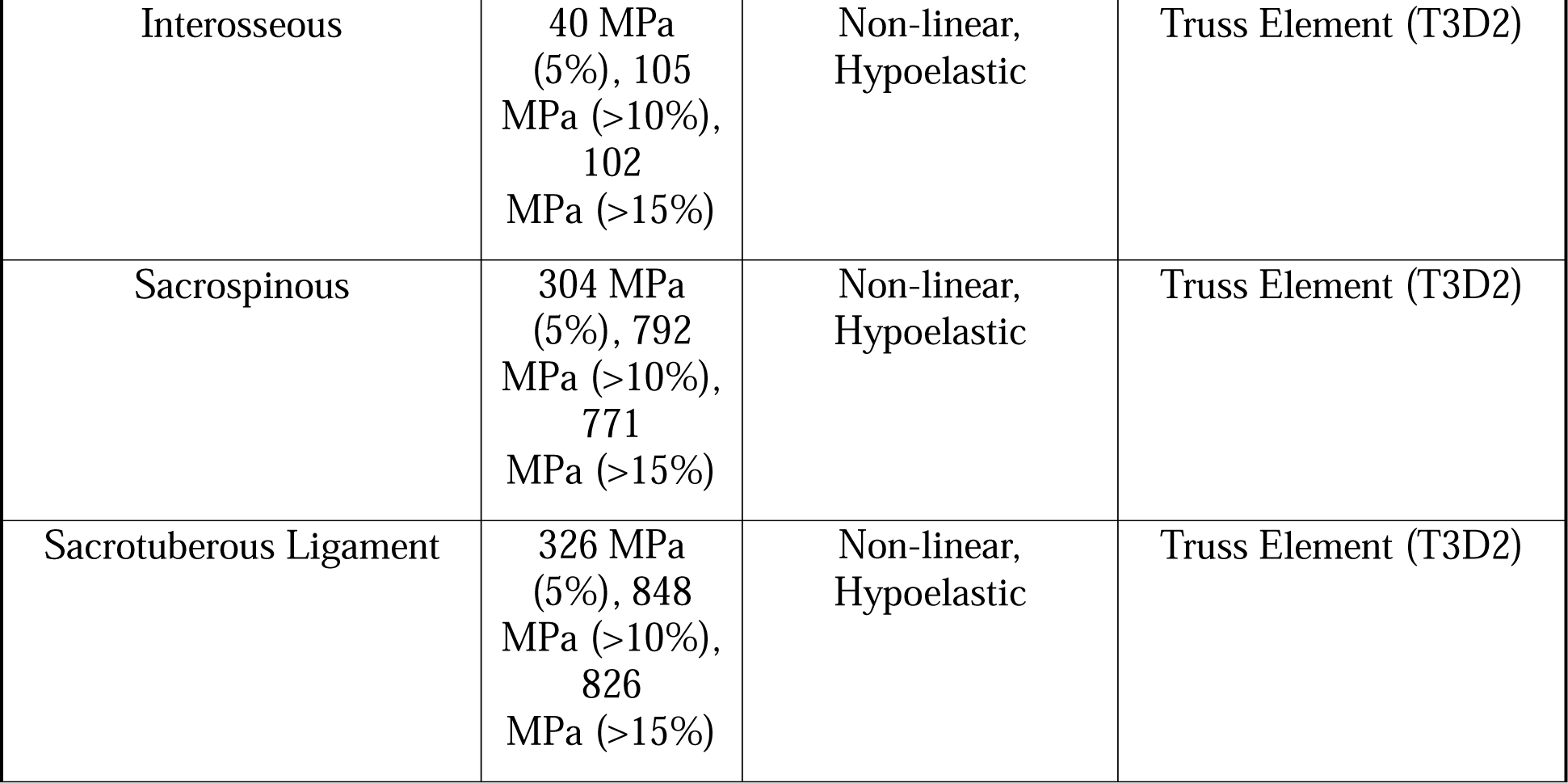
Material properties of the finite element model [17, 41-46]. E represents the Young’s Modulus; v represents Poisson’s Ratio.

| **Component** | **Material Properties** | **Constitute Relation** | **Element Type** |
| --- | --- | --- | --- |
| Vertebral Cortical Bone | E= 12000MPa | Isotropic, Elastic | 8 Node Brick Element (C3D8) |
| | $\nu = 0.3$ | | |
| Vertebral Cancellous Bone | E= 100MPa | Isotropic, Elastic | 4 Node Tetrahedral Element (C3D4) |
| | $\nu = 0.2$ | | |
| Pelvis Cortical Bone (Sacrum, Ilium) | E= 17000MPa | Isotropic, Elastic | 4 Node Tetrahedral Element (C3D4) |
| | $\nu = 0.3$ | | |
| Sacrum Cancellous Bone | Heterogenous | Isotropic, Elastic | 4 Node Tetrahedral Element (C3D4) |
| Ilium Cancellous Bone | E= 70 MPa | Isotropic, Elastic |  |
| | $\nu = 0.2$ | | 4 Node Tetrahedral Element (C3D4) |
| Femur Cortical Bone | E= 17000MPa | Isotropic, Elastic | 4 Node Tetrahedral Element (C3D4) |
| | $\nu = 0.29$ | | |
| Femur Cancellous Bone | E= 100 MPa | Isotropic, Elastic | 4 Node Tetrahedral Element (C3D4) |
| | $\nu = 0.2$ | | |
| Ground Substance of Annulus Fibrosis | $c_{10} = 0.035$ | | |
| | $k_1 = 0.296$ | Anisotropic, Hyperelastic (HGO) | 8 Node Brick Element (C3D8) |
| | $k_2 = 65$ | | |
| Nucleus Pulposus | E= 1 MPa | Isotropic, Elastic | 8 Node Brick Element (C3D8) |
| | $\nu = 0.499$ | | |
| Anterior Longitudinal | 7.8 MPa (<12%), 20 MPa | Non-linear, Hypoelastic | Truss Element (T3D2) |
|  | (>12%) |  |  |
| Posterior Longitudinal | 10 MPa (<11%),<br>20 MPa (>11%) | Non-linear,<br>Hypoelastic | Truss Element (T3D2) |
| Ligamentum Flavum | 15 MPa (<6.2%),<br>19.5 MPa (>6.2%) | Non-linear,<br>Hypoelastic | Truss Element (T3D2) |
| Intertransverse | 10 MPa (<18%),<br>58.7 MPa (>18%) | Non-linear,<br>Hypoelastic | Truss Element (T3D2) |
| Interspinous | 10 MPa (<14%),<br>11.6 MPa (>14%) | Non-linear,<br>Hypoelastic | Truss Element (T3D2) |
| Supraspinous | 8 MPa (<20%), 15 MPa (>20%) | Non-linear,<br>Hypoelastic | Truss Element (T3D2) |
| Capsular | 7.5 MPa (<25%),<br>32.9 MPa (>25%) | Non-linear,<br>Hypoelastic | Truss Element (T3D2) |
| Anterior SIJ | 125 MPa (5%), 325 MPa (>10%),<br>316 MPa (>15%) | Non-linear,<br>Hypoelastic | Truss Element (T3D2) |
| Short Posterior SI | 43 MPa (5%), 113 MPa (>10%),<br>110 MPa (>15%) | Non-linear,<br>Hypoelastic | Truss Element (T3D2) |
| Long Posterior SI | 150 MPa (5%), 391 MPa (>10%),<br>381 MPa (>15%) | Non-linear,<br>Hypoelastic | Truss Element (T3D2) |
| Interosseous | 40 MPa (5%), 105 MPa (>10%), 102 MPa (>15%) | Non-linear, Hypoelastic | Truss Element (T3D2) |
| Sacrospinous | 304 MPa (5%), 792 MPa (>10%), 771 MPa (>15%) | Non-linear, Hypoelastic | Truss Element (T3D2) |
| Sacrotuberous Ligament | 326 MPa (5%), 848 MPa (>10%), 826 MPa (>15%) | Non-linear, Hypoelastic | Truss Element (T3D2) |

### Finite Element Model Validation

Previous *in-vitro* studies were used to validate the current FE model’s spinal range of motion (ROM) for both the thoracic and lumbar segments [15, 16]. The thoracic (T1-T12) ROM of the finite element model was compared to *in-vitro* ROM data generated by Watkins et al [16]. Lumbar (L1-S1) ROM was validated by comparing *in-vitro* ROM from Panjabi et al [15]. This model was subject to a 4 Nm at the T1 vertebral body [8, 14] and at 6 Nm at the L1 vertebral body to obtain range of motion (ROM) for both spinal flexion and extension [8, 17].

### Finite Element Surgical Models

Three models were created from this original model to simulate the traditional laminectomy and bilateral and unilateral midline sparing approaches at the L4-L5 segment. An L4-L5 laminectomy was simulated by removing the spinous process, supraspinous, and interspinous ligaments. Bony elements of the L4 lamina from the L4 pedicle to the L5 pedicle were resected. Additionally, the ligamentum flavum was removed and the medial 1/3th of the right and left L4-L5 facet joint was resected [2]. Similarly, the bilateral laminotomies were modelled by resecting the bone from the inferior 1/3 of the L4 lamina and the superior 1/3 of the L5 lamina on the right and left sides, whereas the unilateral laminotomy was simulated on the right side only. The spinous process, supraspinous, and interspinous ligaments, pars interarticularis and fact capsule were preserved in these laminotomy cases (Figure 1) [2]. The same moment was applied to these models and segmental ROM was obtained for flexion and extension.

**Figure 1:**
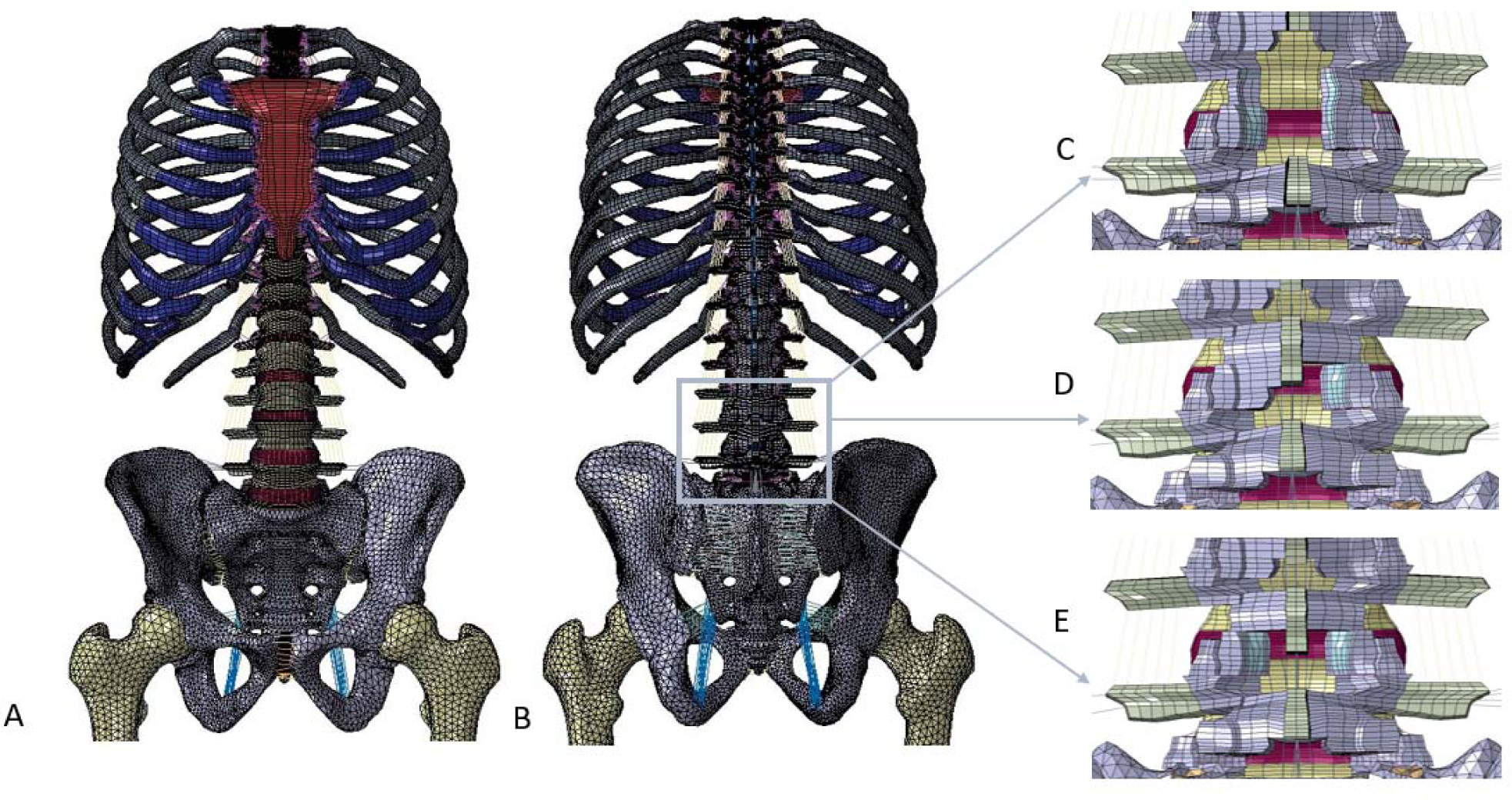
Finite element model. A represents anterior view of the model with no surgery. B represents the posterior view of the model with no surgery. C represents the posterior view of the L4-L5 laminectomy model. D represents the posterior view of the unilateral L4-L5 laminotomy case and E represents the posterior view of the bilateral L4-L5 laminotomy.

### Musculoskeletal Models

The segmental ROM for all models were input into the OpenSim (SimBios, Stanford University, Stanford, CA) thoracolumbar musculoskeletal model constructed and validated by Bruno et al [18] (Figure 2). In short, these models have 93 degrees-of-freedom and 552 musculotendon actuators [18].As discussed in the literature, the laminotomy procedure involves 18.7% CSA damage to the procedure site [1]. Therefore, the OpenSim model for bilateral and unilateral laminotomy was adjusted by reduction of the maximum isometric force [F^M^_max_ (N) = CSA (cm^2^)* specific tension (N/cm^2^)] in the multifidus (MF) muscles. The laminectomy procedure involved complete damage of the MF fibers at the surgical site. Muscle force estimation was performed by utilizing the static optimization tool in OpenSim, which estimates muscle activations and forces by distributing net joint moments across individual muscle forces at successive time point. This process involves minimizing the sum of squared muscle activations. Following static optimization within this model, the output was the static optimization force.sto file, from which we derived the time histories of muscle forces.

**Figure 2:**
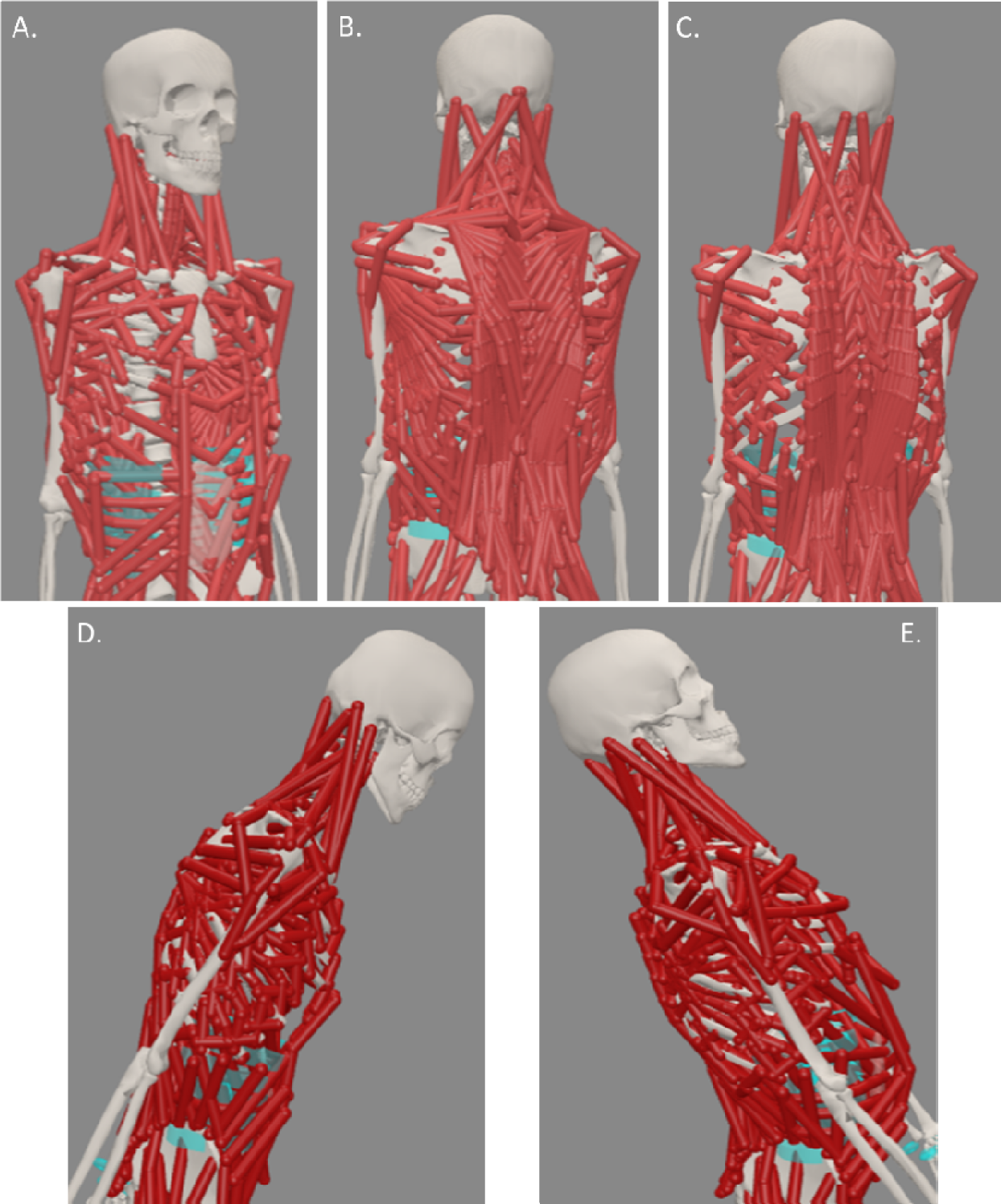
OpenSim Thoracolumbar model [18] showing the [A.] anterior, [B.] posterior with all muscles, and [C] posterior with the superficial muscles (trapezius, latissimus dorsi) removed. This model was created and validated by Bruno AG et. al. [4]. ROM obtained from 10 Nm moment was applied to the model in [D] flexion and [E] extension.

### Data Analysis

Muscle forces were obtained and analyzed to determine the amount of usage with extension and flexion. The muscles included the latissimus dorsi (LD), trapezius (trap), longissimus thoracis pars lumborum (LTpL), longissimus thoracis pars thoracis (LTpT), multifidus (MF), Iliocostalis Lumborum (IL), Psoas Major (PM), Quadratus Laborum (QL), Transversus Abdominus (TA), Erector Spine (ES), External Oblique (EO), and Internal Oblique (IO), Internal Intercostal (II), and External Intercostal (EI). We first took the percent change of muscle tension when comparing a unilateral to bilateral laminotomy. We then took the percent change of each muscle force to determine how the muscle use changed when compared to a normal spine. We also compared the percent change when comparing a laminotomy to a laminectomy. We further investigated the muscle forces that cross over each vertebral disc level. These muscle forces allowed us to analyze the amount of inner disk pressure (IDP) experienced at each vertebral level.

## Results

### Range of motion

The ROM of each case fell within range of the *in-vitro* data found by Panjabi et al and Watkins et al [15, 16]. This data can be found in the supplemental data section.

### Unilateral vs Bilateral Laminotomy

Muscle forces when comparing bilateral and unilateral laminotomy displayed a percent change of less than 5% for every muscle group in both flexion and extension (See figure 3). The exception to this was the TA displayed a 13% change in force favoring the bilateral laminotomy when the spine is in flexion. Furthermore, IL and QL saw a 5.3% and 8.3% change favoring unilateral laminotomy when in spinal extension. When the forces displayed over each vertebral level were calculated, there was no difference in percent change higher than 5%. Due to the two procedures having similar muscle force changes as well as tension changes over levels of the spine, a bilateral laminotomy was used for the rest of the study.

**Figure 3:**
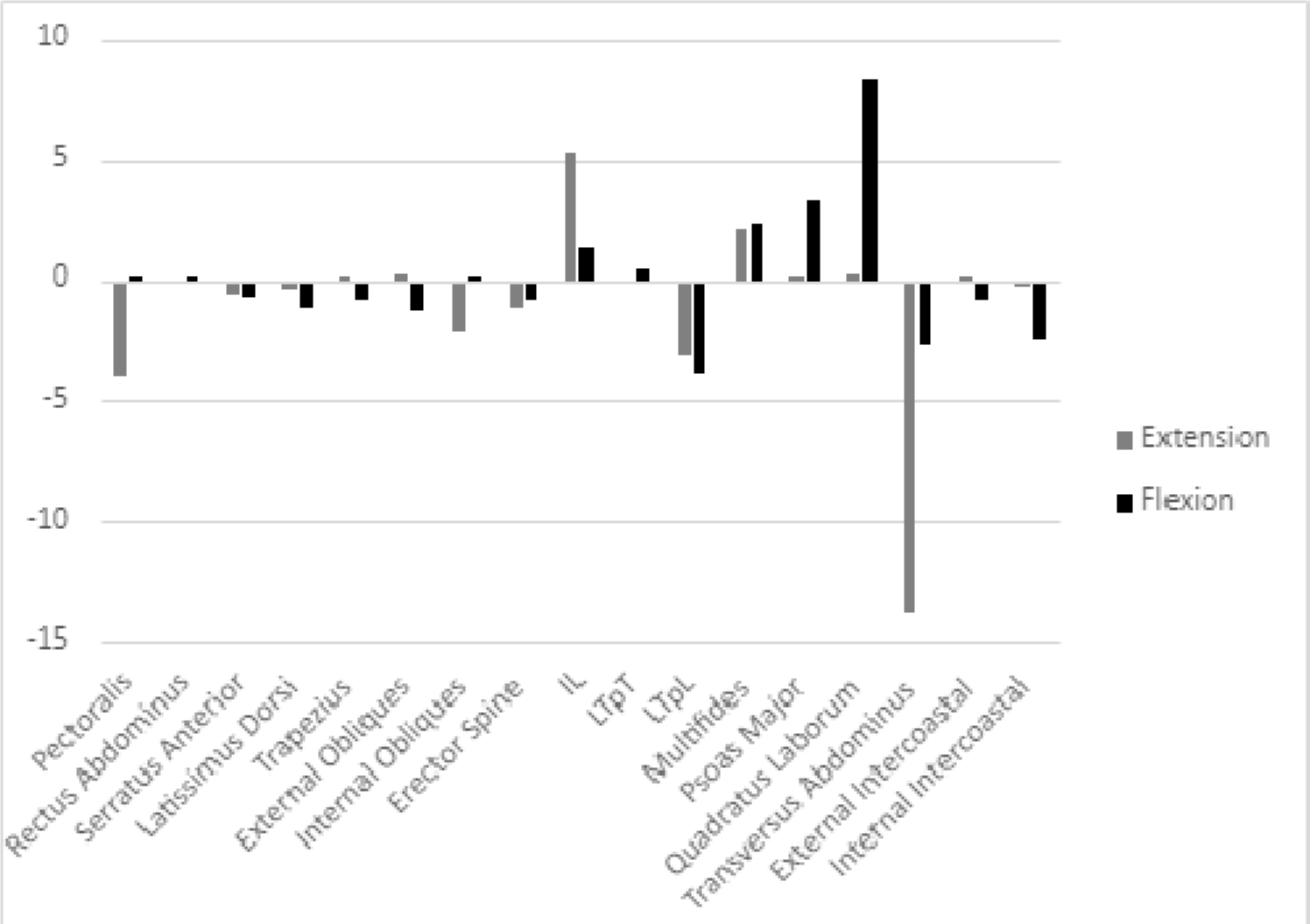
Percent change of muscle force when comparing the bilateral and unilateral laminotomy. Above the x-axis favors unilateral laminotomy and below favors bilateral laminotomy.

### Laminotomy vs Normal Spine

When looking at the effect of a laminotomy on the muscles of the back, there was an increase in force in the IL by 12% and decrease in TA by 25% when the spine is in extension. All other muscles did not have a change greater than 10%. When the spine was flexed all muscles displayed less than a 10% change in force (See figure 4a).

**Figure 4:**
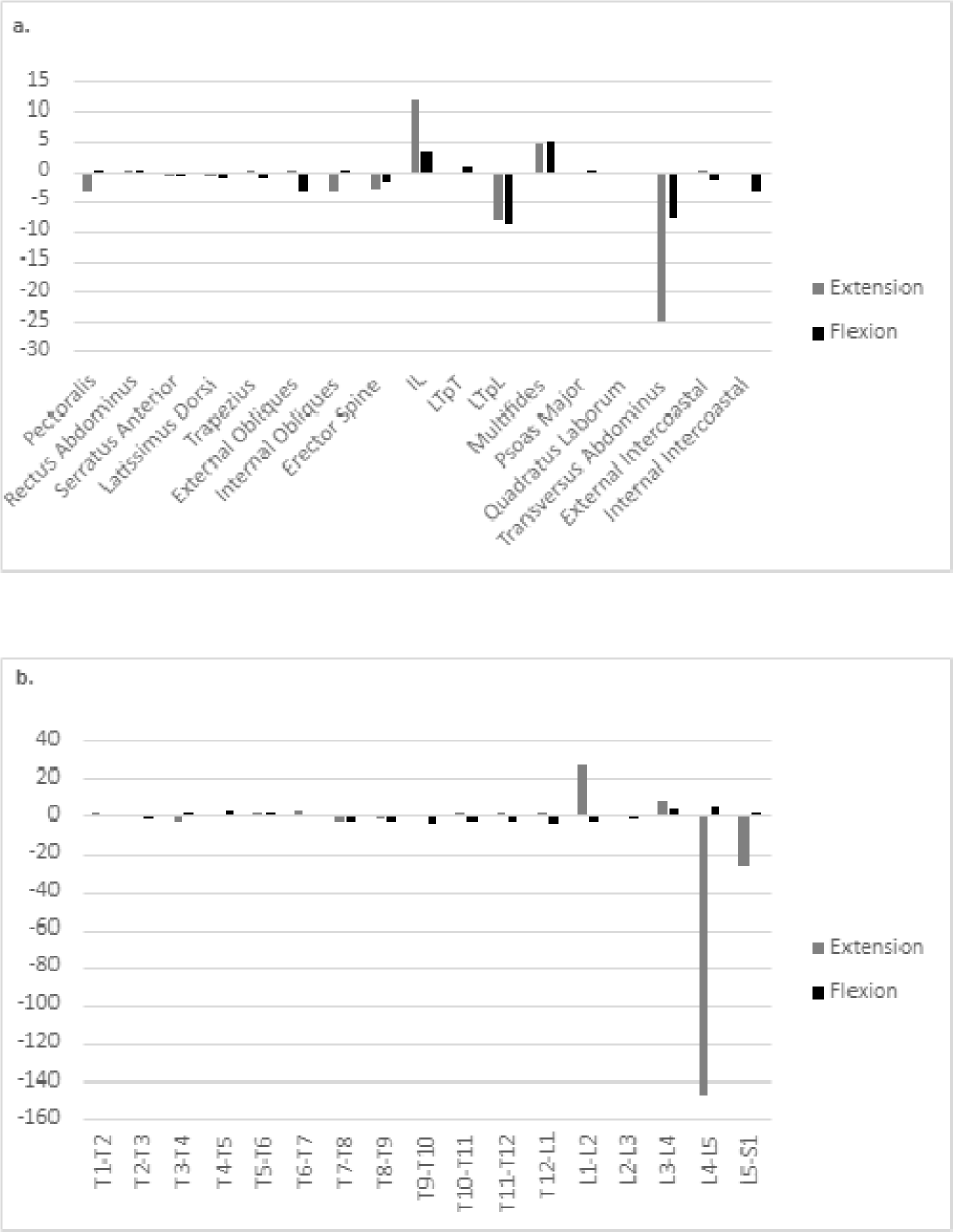
Demonstrates percent change when comparing the normal spine with a spine that has undergone a laminotomy in both flexion and extension. [a.] demonstrates the percent changes of the muscular tension forces while [b.] demonstrates the percent change of the stresses over each level of the spine. Values above the x-axis favor the spine with no interventions while below the x-axis favors a spine that has undergone a laminotomy.

When the forces over vertebral levels was analyzed, there was no percent difference between an intact spine and laminotomy greater than 10% when the spine was in extension. However, when the spine was in flexion, there was an increase in force by 147% across L4-L5 and 27% across L5-S1. There was also a decrease in force across L1-L2 by 27% (See figure 4b).

### Laminectomy vs Normal Spine

When studying the effects of a laminectomy in muscle forces, there was an increase in IO by 35%, LTpL by 10%, and TA by 255% when the spine was in extension. There was also a decrease in IL by 23% and 20% in MF. When the spine was placed in flexion there was increase tension in LD by 65%, Trap by 13%, EO by 13%, IO by 59%, LTpL by 57%, and II by 40%. There was also a decrease in SA by 14%, IL by 37%, and MF by 16% (See figure 5a).

**Figure 5:**
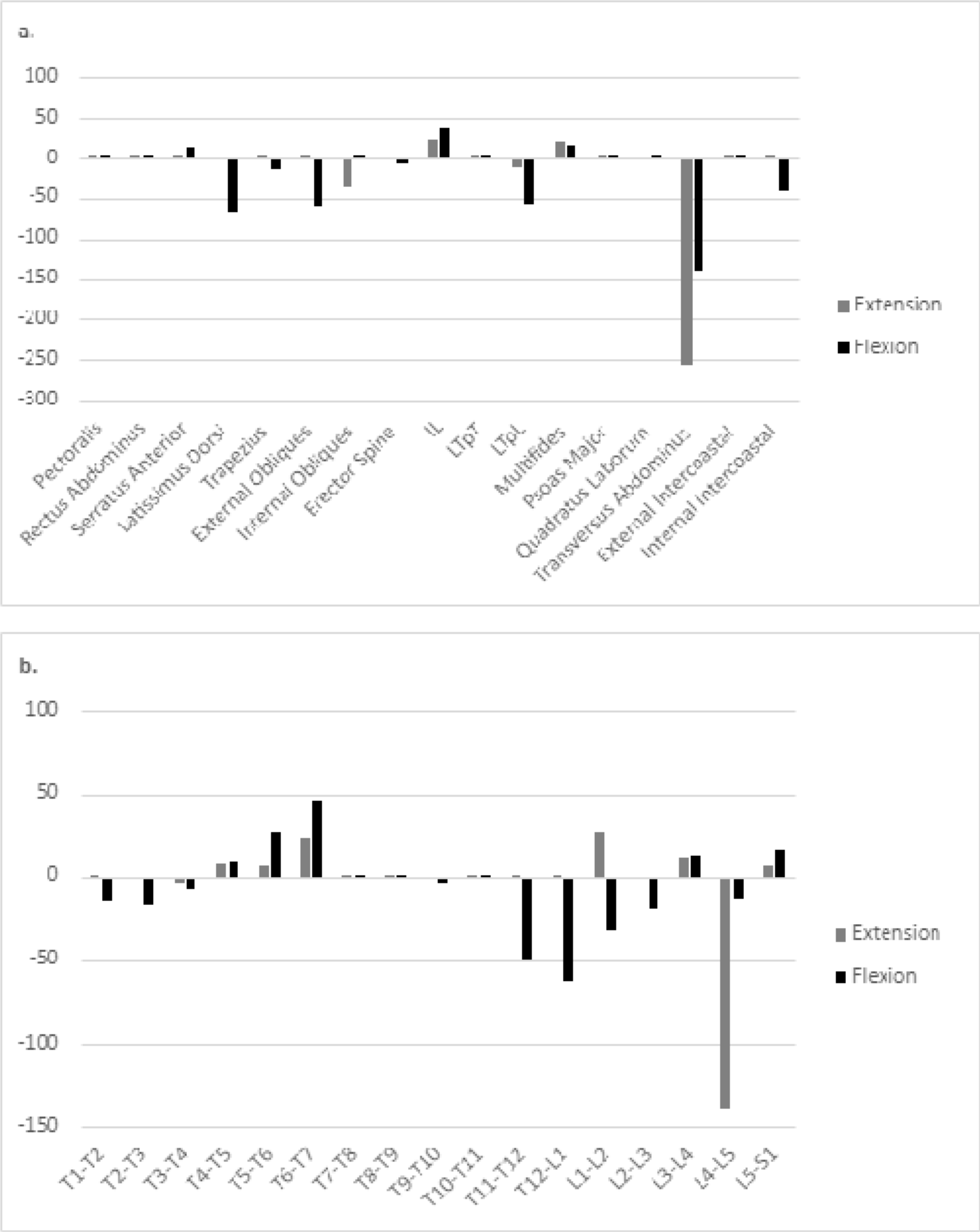
Demonstrates percent change when comparing the normal spine with a spine that has undergone a laminectomy in both flexion and extension. [a.] demonstrates the percent changes of the muscular forces while [b.] demonstrates the percent change of the stresses over each level of the spine. Values above the x-axis favor the spine with no interventions while below the x-axis favors a spine that has undergone a laminectomy.

The force present over vertebral levels when the spine was in extension increased in T1-T2 by 13%, T2-T3 by 17%, T11-T12 by 50%, T12-L1 by 62%, L1-L2 by 32%, L2-L3 by 18%, L4-L5 by 12%. There was also a decrease in force over T5-T6 by 26%, T6-T7 by 46%, L3-L4 by 13%, and L5-S1 by 16%. When the spine was placed in flexion there was an increase in force over L4-L5 by 139%. There was also a decrease in force over T6-T7 by 23%, L1-L2 by 27%, and L3-L4 by 13% (See figure 5b).

### Laminotomy vs Laminectomy

Comparing the muscle tension in laminotomy to the laminectomy when the spine was in extension displayed an increase in IL by 12% and multifidus by 16%. There was also a decrease in tension within the TA by 138% and ES by 12%. When placed in flexion there was an increase in SA by 15%, IL by 35%, and MF by 12%. There was a decrease in LD by 63%, Trap by 11% and EO by 54%, LTpL by 44%, and TA by 123%. There was also an increase in tension within the SA by 15%, IL by 35%, and MF by 12% (see figure 6a).

**Figure 6:**
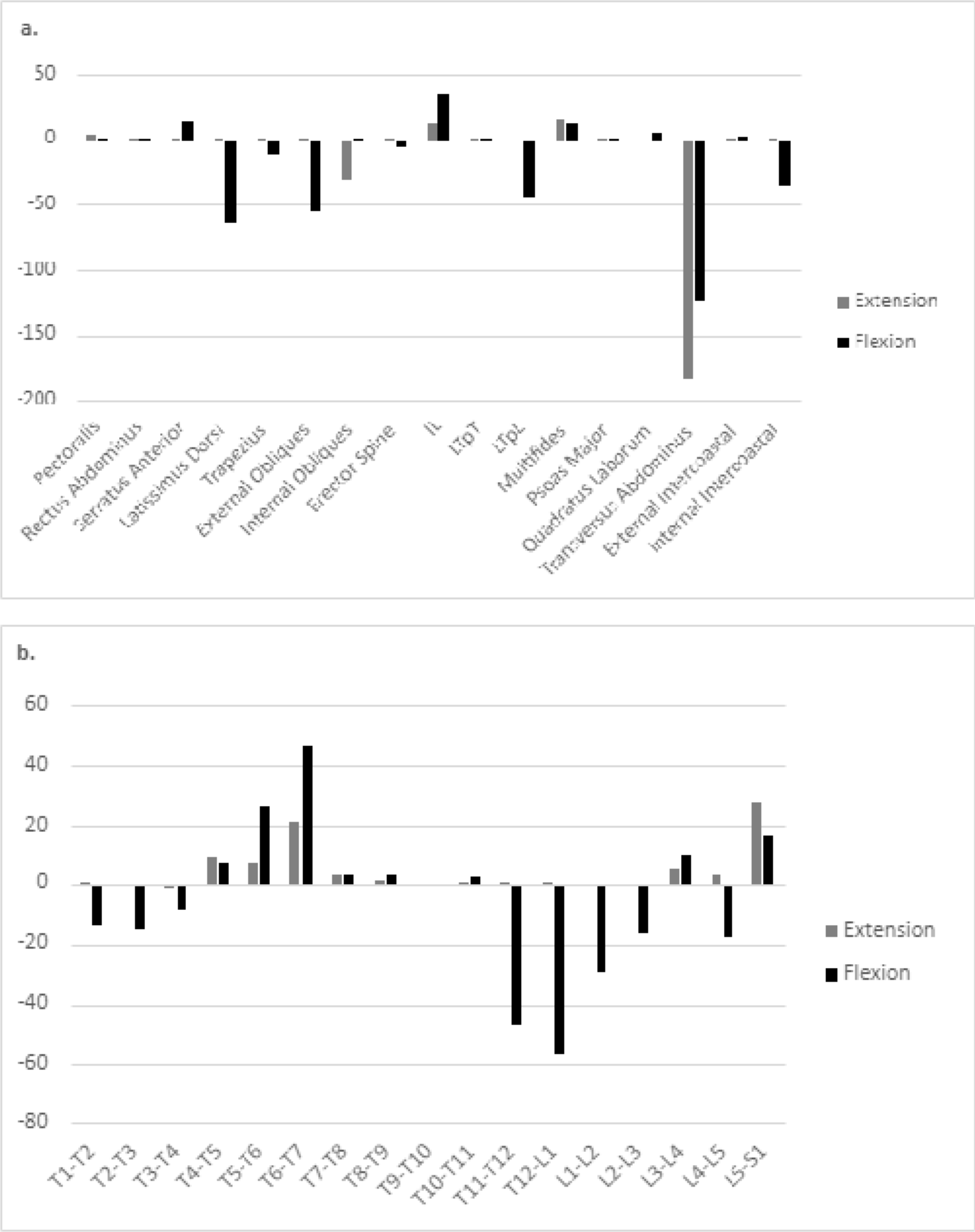
Demonstrates percent change when comparing a spine that has undergone a laminotomy with one that has undergone a laminectomy in both flexion and extension. [a.] demonstrates the percent changes of the muscular forces while [b.] demonstrates the percent change of the stresses over each level of the spine. Values above the x-axis favor the spine after a laminotomy while below the x-axis favors a spine that has undergone a laminectomy.

The forces across levels when looking at the spine in extension displayed in increase by T5-T6 by 26%, T6-T7 by 47%, and L5-S1 by 16%. There was also a decrease in forces across in T1-T2 by 13%, T2-T3 by 15%, T11-T12 by 47%, T12-L1 by 56%, L1-L2 by 29%, L2-L3 by 17%, L4-L5 by 17%. When the spine was placed in flexion there was an increase across T6-T7 by 21% and L5-S1 by 28% (See figure 6b).

## Discussion

The data collected though our simulations suggest that laminotomies, by preserving the multifidus (MF) group, results in less muscular compensation compared to laminectomies. The preservation of the MF muscles enables a post-laminotomy spine to function more similar to the normal spine, in contrast to the post-laminectomy spine. Additionally, a large muscle load transfer was seen in the thoracic segments (T4-T7) following the laminectomy compared to a normal spine, with reduced flexion muscle forces at the thoracolumbar transition zone (T11-L2). Furthermore, the muscle group which showed compensatory action among all cases was the erector spinae (ES) (IL) group and undamaged MF fibers. Our findings suggest that the minimally invasive conservation of muscle integrity in laminotomy procedures could be crucial for maintaining closer-to-normal spinal function, highlighting the potential benefits of this surgical approach in reducing the need for muscular compensation.

A biomechanical study by Ho et al [19] found comparable results when assessing flexion and extension in cadaver specimens that underwent unilateral and bilateral laminotomies, agreeing with our results which showed minimal intersegmental muscle changes among the laminotomy cases. However, our results also indicated that muscles attaching across the L5-S1 segment also reduced in load, with increases at the L1-L2 level. Previous FE analyses of the spine simulating the laminectomy procedure showed significant instability when removing soft tissue elements which stabilize the spine [20–22]. Conversely, studies that have compared a laminectomy to laminotomy reveal the latter maintains greater stability in the lumbar region [19, 23, 24]. An increase in spinal instability will result in increased muscle compensation to stabilize the lumbar spine. In fact, studies have shown that contraction of the paraspinal extensor muscles such as the MF provide stability against shear forces in during loading of the lumbar spine [25]. Additionally, research findings suggest that it is essential to preserve the posterior ligamentous complex during these lumbar operations. A previous biomechanical study by Tai et al emphasized that preserving the posterior tension band results in a decreased likelihood of developing instability in the lumbar spine [26]. Preservation of the bones and ligaments of the spine results in greater preservation of normal motion of the lumbar spine after surgery [27]. Our results indicates that laminectomies, by reducing the stabilizing function of the MF muscles, necessitate increased muscular compensation to restore lost stability. This compensation emphasizes the critical role of MF muscles in spinal stability and the large demand placed on remaining muscles to maintain spinal integrity.

Destruction of the posterior band also plays a significant role in the stress experience between adjacent vertebral levels, emphasizing the protective role of paraspinal muscles against segment loading. Damage to these paraspinal muscles in open procedures such as laminectomies, have been shown to increase the shear forces between at the rostral and caudal levels [28]. An FE study conducted by Kumaran et al showed that after damaging the MF fibers in open surgery spinal fusion, stresses were increased at the adjacent segment intervertebral discs [8]. Our results agree with these findings as the open surgery of the laminectomy displayed an increase in muscle forces at the rostral and caudal adjacent segments compared to the laminotomy cases. Our study may provide evidence that minimally invasive surgeries on the spine decrease spinal element stresses and potentially decrease post-operative adjacent segment degeneration. This may provide evidence that the MF are crucial in preserving the force between vertebral levels, especially in the lumbar region.

Clinically, research has shown that patients who have undergone surgical decompression for spinal stenosis have a greater improvement in pain and function [29]. Both an open and minimally invasive approach are effective methods to relieve the symptoms associated with lumbar stenosis [30]. Previous studies revealed that there were no clinical differences in preoperative symptom improvement when groups received a unilateral laminotomy compared to those receiving a laminectomy [5]. However, research has shown that there is significant post-operative spinal instability in rotation and flexion following the laminectomy procedure [31, 32]. Furthermore, patients who receive a laminotomy had advantages in recovery, postoperative stability, and postoperative rehabilitation outcomes [33]. Additional clinical studies have shown similar results for the treatment of bilateral spinal stenosis with the unilateral and bilateral approach. Overall patient satisfaction and the reduction of symptoms is similar when comparing these two laminotomy procedures [34, 35]. At a 10 year follow up, decompression via laminotomy displayed effective reduction in symptoms [36]. Further studies have illustrated that using a unilateral laminotomy is also an effective procedure for a bilateral decompression [37]. This shows achievement in short-term clinical results for spinal stenosis, regardless of the surgical technique utilized. However, a previous clinical study has shown that increased retraction of the paraspinal muscles of the spine lead to worse functional outcomes and increased pain [6]. This study showed that increased intramuscular pressure overall leads to a higher degree of backpain. We found that there are increases in muscle forces in the laminectomy case which may be associated with increased pain in a similar way that the intramuscular pressure produced more post operative pain.

As with any computational study, there are limitations which should be mentioned. FE simulations were used to generate the spinal kinematic data used to drive the musculoskeletal model to generate the muscle force data. Despite the lack of *in-vivo* kinematic data, our spine model’s kinematics have been rigorously validated with several studies to ensure proper motion in all motions[8, 10, 11, 15, 38]. Only static optimization (SO) was used to estimate muscle activations. Computed muscle control (CMC) is another method to compute muscle excitations to drive musculoskeletal models. The static optimization technique was used in this study because some studies had suggested that SO was the superior optimization technique for estimating muscle function due to its robustness and computational efficiency [39, 40].

## Conclusion

Our study highlights the biomechanical impact of surgical approach on spinal stability and muscle compensation in spinal stenosis treatment. Our results highlight that laminectomy, which involves the removal of paraspinal muscles and posterior ligamentous structures to relieve stenosis, can lead to increased instability and necessitate muscle compensation, particularly in adjacent and thoracic spine segments. Conversely, midline sparing approaches such as laminotomies, are associated with decreased muscle compensation across spinal segments and enhanced stability. These findings advocate for the preservation of paraspinal musculature to maintain spinal integrity. Future research is essential to further validate these conclusions and guide surgical decision-making in treating lumbar spinal stenosis.

## Supplemental Data

**Supplement Table 1:**
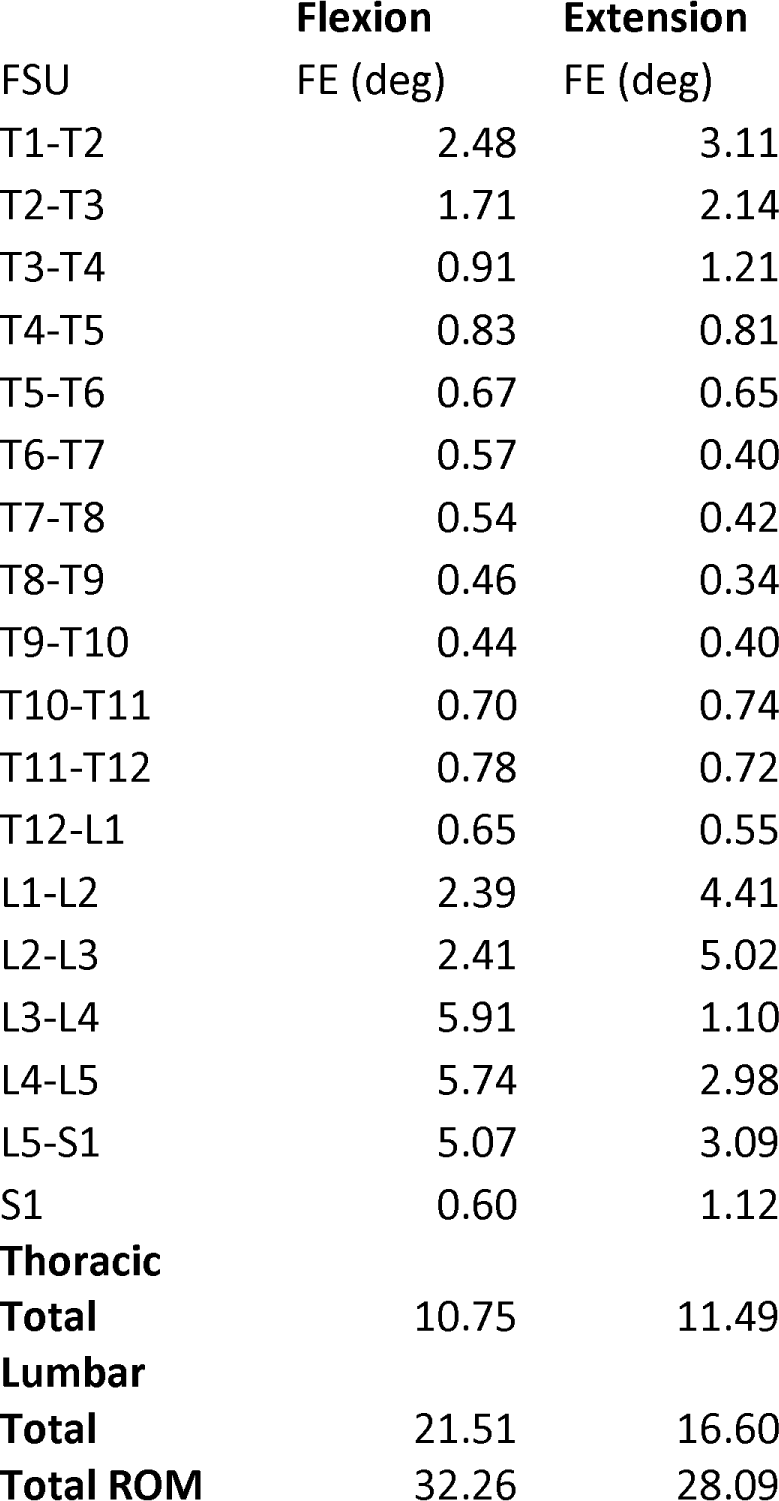
ROM of the laminectomy model in flexion and extension [15, 16].

|  | **Flexion** | **Extension** |
| --- | --- | --- |
| FSU | FE (deg) | FE (deg) |
| T1-T2 | 2.48 | 3.11 |
| T2-T3 | 1.71 | 2.14 |
| T3-T4 | 0.91 | 1.21 |
| T4-T5 | 0.83 | 0.81 |
| T5-T6 | 0.67 | 0.65 |
| T6-T7 | 0.57 | 0.40 |
| T7-T8 | 0.54 | 0.42 |
| T8-T9 | 0.46 | 0.34 |
| T9-T10 | 0.44 | 0.40 |
| T10-T11 | 0.70 | 0.74 |
| T11-T12 | 0.78 | 0.72 |
| T12-L1 | 0.65 | 0.55 |
| L1-L2 | 2.39 | 4.41 |
| L2-L3 | 2.41 | 5.02 |
| L3-L4 | 5.91 | 1.10 |
| L4-L5 | 5.74 | 2.98 |
| L5-S1 | 5.07 | 3.09 |
| S1 | 0.60 | 1.12 |
| **Thoracic** |  |  |
| **Total** | 10.75 | 11.49 |
| **Lumbar** |  |  |
| **Total** | 21.51 | 16.60 |
| **Total ROM** | 32.26 | 28.09 |

**Supplement Table 2:** ROM of the Unilateral Laminotomy model in flexion and extension [15, 16].

|  | **Flexion** | **Extension** |
| --- | --- | --- |
| FSU | FE (deg) | FE (deg) |
| T1-T2 | 3.99 | 3.11 |
| T2-T3 | 2.58 | 2.14 |
| T3-T4 | 1.21 | 1.21 |
| T4-T5 | 0.86 | 0.81 |
| T5-T6 | 0.50 | 0.64 |
| T6-T7 | 0.30 | 0.40 |
| T7-T8 | 0.26 | 0.42 |
| T8-T9 | 0.20 | 0.34 |
| T9-T10 | 0.16 | 0.40 |
| T10-T11 | 0.20 | 0.74 |
| T11-T12 | 0.26 | 0.72 |
| T12-L1 | 0.22 | 0.55 |
| L1-L2 | 3.13 | 4.61 |
| L2-L3 | 2.87 | 5.22 |
| L3-L4 | 3.96 | 1.21 |
| L4-L5 | 4.89 | 2.18 |
| L5-S1 | 6.52 | 3.31 |
| S1 | 0.73 | 1.19 |
| **Thoracic** |  |  |
| **Total** | 10.75 | 11.49 |
| **Lumbar** |  |  |
| **Total** | 21.38 | 16.53 |
| **Total ROM** | 32.13 | 28.02 |

**Supplemental Table 3:** ROM of the bilateral laminotomy model in flexion and extension [15, 16].

|  | **Flexion** | **Extension** |
| --- | --- | --- |
| FSU | FE (deg) | FE (deg) |
| T1-T2 | 3.99 | -3.11 |
| T2-T3 | 2.58 | -2.14 |
| T3-T4 | 1.21 | -1.21 |
| T4-T5 | 0.86 | -0.81 |
| T5-T6 | 0.49 | -0.64 |
| T6-T7 | 0.31 | -0.40 |
| T7-T8 | 0.26 | -0.42 |
| T8-T9 | 0.20 | -0.34 |
| T9-T10 | 0.16 | -0.40 |
| T10-T11 | 0.20 | -0.74 |
| T11-T12 | 0.26 | -0.72 |
| T12-L1 | 0.22 | -0.55 |
| L1-L2 | 3.13 | -4.56 |
| L2-L3 | 2.87 | -5.18 |
| L3-L4 | 3.95 | -1.19 |
| L4-L5 | 4.92 | -2.38 |
| L5-S1 | 6.51 | -3.25 |
| S1 | 0.73 | -1.17 |
| **Thoracic** |  |  |
| **Total** | 10.75 | 11.49 |
| **Lumbar** |  |  |
| **Total** | 21.38 | 16.55 |
| **Total ROM** | 32.13 | 28.04 |

